# Isolation and electrochemical analysis of a facultative anaerobic electrogenic strain *Klebsiella* sp. SQ-1

**DOI:** 10.1101/2023.12.02.569720

**Authors:** Lei Zhou, Tuoxian Tang, Dandan Deng, Yayue Wang, Dongli Pei

**Affiliations:** Henan Provincial Engineering Research Center for Development and Application of Characteristic Microorganism Resources, College of Biology and Food, Shangqiu Normal University, Shangqiu, Henan 476000, PR China; Department of Biological Sciences, Virginia Tech, Blacksburg, VA 24060, USA

**Keywords:** microbial fuel cell, *Klebsiella* sp., electrochemical analysis, extracellular respiration, facultative electricigens

## Abstract

Electricigens decompose organic matter and convert stored chemical energy into electrical energy through extracellular electron transfer. They serve as significant biocatalysts for microbial fuel cells which have practical applications in green energy generation, effluent treatment, and bioremediation. A facultative anaerobic electrogenic strain SQ-1 is isolated from sludge in a biotechnology factory. The strain SQ-1 is a close relative of *Klebsiella variicola*. Multilayered biofilms form on the surface of a carbon electrode after the isolated bacteria are inoculated into a microbial fuel cell device. This strain produces high current densities of 625 μA cm^-2^ by using acetate as the carbon source in a three-electrode configuration. The electricity generation performance is also analyzed in a dual-chamber microbial fuel cell. It reaches a maximum power density of 560 mW m^-2^ when the corresponding output voltage is 0.59 V. The facultative strain SQ-1 utilizes hydrous ferric oxide as an electron acceptor to perform extracellular electricigenic respiration in anaerobic conditions. Since facultative strains possess better properties than anaerobic strains, *Klebsiella* sp. SQ-1 may be a promising exoelectrogenic strain for applications in microbial electrochemistry.

## Introduction

Microbial fuel cell (MFC) is a bio-electrochemical system that converts chemical energy to electricity by using electricigens as biocatalysts, which has practical values in biomass degradation, sewage disposal, and clean energy generation (Elmaadawy et al., 2022; Ren et al., 2022; Thapa et al., 2022). Electricigens function as biocatalysts in MFC, which play vital roles in substrate oxidation and electron (or hydrogen) transfer. The electricity generation process consists of several steps: anodic oxidation, electron transport in an external circuit, proton migration, and cathodic reduction. Generally, two types of inocula are utilized in MFC: mixed culture and pure culture. In mixed culture, microbial interactions offer good adaptability to the environment because the microflora can degrade a wide variety of substrates and facilitate concurrent power generation. Additionally, some microbes secrete specific electron transfer mediators (ETMs) which may also contribute to the performance boost in the MFC (Islam et al., 2018; Kiely et al., 2011). However, this synergy effect in microflora doesn’t always happen. Antagonistic interactions and metabolic conflicts between different microbes give rise to a performance decrease in MFC.

In contrast, using simple substrates as carbon sources such as acetate, formate, citrate, and pyruvate (Yousaf, 2017; Zhao et al., 2017; Zhou et al., 2016), pure culture achieved better performance than mixed culture. Different types of electricigens like *Geobacter* sp., *Shewanella* sp., *Pseudomonas* sp., *Citrobacter* sp., *Klebsiella* sp., *Escherichia* sp., *Bacillus* sp., and *Saccharomyces cerevisiae* have been employed as biocatalysts in MFC (Bond et al., 2003; Cao et al., 2019; Ieropoulos et al., 2005; Jimenez Pacheco et al., 2023; Kim et al., 2005; Rabaey et al., 2005). Studies on these isolated bacteria helped scientists better understand the specific mechanisms of extracellular respiration (Bishir et al., 2023; Meylani et al., 2023). More importantly, it is easier to manipulate these electricigens and optimize their power generation performance in MFC. Among the studied electricigens, *Geobacter* sp. and *Shewanella* sp. are the most representative anaerobic and facultative anaerobic strains, respectively. *Geobacter* sp. can use ambient ferric iron or solid electrodes as electron acceptors for extracellular electron transfer, but it is not able to secret ETMs. As for *Shewanella* sp., the “nanowires” extended from the outer membrane give it an advantage in extracellular electron transfer, by which the ferric reduction is accelerated. However, it showed much lower power generation in MFC or other bio-electrochemical systems than strictly anaerobic *Geobacter* sp. under the same conditions (Marsili et al., 2008; Pirbadian et al., 2014).

To date, several facultative anaerobic electricigens such as *Shewanella*sp., *Pseudomonas* sp., *Citrobacter* sp., and *Klebsiella* sp. were isolated and able to achieve the metabolic pathway shift between electricigenic respiration and anaerobic fermentation in MFC, which showed prospective applications in bioremediation, wastewater treatment, heavy metal leaching, and the global carbon cycle (Chaudhuri et al., 2003; Deng et al., 2010; Rabaey et al., 2004). For instance, the wild-type *Klebsiella variicola* generates electricity by utilizing the palm oil mill effluent as a substrate, achieving the maximum power density of 4,426 mW m^-3^ and a high coulombic efficiency of 63% in MFC (Islam et al., 2018). Another facultative anaerobic electricigen named *Shewanellaoneidensis* MR-1 is a model dissimilatory iron-reducing bacterium, which uses iron oxides as electron acceptors and adsorbs phosphate anions and metal ions during the bioreduction process (Ge et al., 2022; Long et al., 2021; Yu et al., 2022).

Previous electrochemical studies have demonstrated that the low power output by electricigens is a big hurdle to the industrial applications of MFC (Bello et al., 2017). Therefore, it is a long-term challenge to screen superior exoelectrogenic strains with high efficiency and strong environmental adaptability. In this study, a facultative anaerobic strain SQ-1 was isolated from sludge in a biotechnology factory. The electricity generation capacity and ferric reduction of this strain were investigated by using dual-chamber MFCs and an electrochemical workstation.

## Experimental

### Materials and methods

#### Growth medium, inoculation, and culture

The activated sludge was collected from an anaerobic waste water tank in New Yangshao Biological Technology Factory, Henan, China. The sample was inoculated in 100 mL of growth medium containing 0.6 g KCl, 1.5 g NH_4_Cl, 0.3 g KH_2_PO_4_, 0.1 g MgCl_2_, 0.1 g CaCl_2_, 10 mL of trace element solution, and 10 mL of vitamin solution as previously described (Liu et al., 2014; Zhou et al., 2016). Meanwhile, 20 mM acetate sodium and 40 mM fumarate were supplemented as electron donors and acceptors, respectively. The initial pH of the medium was adjusted to 7.2 and autoclaved at 121 °C for 15 min. The mixed microflora (or pure bacteria) was incubated at 30 °C under aerobic or anaerobic static conditions. Anaerobic conditions were created in serum vials by purging with high-purity nitrogen for 20 min and the vial was sealed with a rubber plug. The model electricigen *G. sulfurreducens* PCA was purchased from Deutsche Sammlung von Mikroorganismen und Zellkulturen GmbH and cultured in the growth medium.

#### Isolation and identification of the electricigenic strain

The enrichment of facultative anaerobes was achieved by injecting 10% of the sludge sample into 200 mL of growth medium in a 500 mL flask. The microbes were cultured at 30 °C, 180 rpm for 7 days. Five percent of the selective inoculation was transferred into the growth medium under anaerobic conditions and cultured at 30 °C for 7 days to further deplete aerobic bacteria. The finally as-obtained mixed bacteria were utilized as inoculum for screening facultative anaerobic electricigens using a dual-chamber MFC.

After the electrochemically active biofilm was enriched on the anode, the whole electrode was immersed in the phosphate-buffered saline (PBS) solution and the biofilm was detached by shaking strongly to prepare a bacterial suspension for utilization. The well-dispersed samples were serially diluted from 10^-1^ to 10^-6^ and plated on agar plates in an anaerobic incubator. After one week of culture at 30 °C, different colonies were inoculated into the liquid growth medium. An MFC device was used to test voltage-producing capabilities. The whole isolation procedure was repeated until single colonies of facultative anaerobic electricigens were isolated and they exhibited similar output power density.

Discrete colonies were used as templates for PCR. The following universal primers were used to amplify the 16S rDNA: 27F (5′-AGA GTTTGA TCC TGG CTC AG-3′) and 1492R (5′-GGT TAC CTT GTT ACGACT T-3′). The 16S rDNA gene sequences were compared in the National Centre for Biotechnology Information (NCBI) databank, and the related neighbor-joining phylogenetic tree was constructed by using the Molecular Evolutionary Genetics Analysis (MEGA, version 6.0) package. A bootstrap analysis was based on 1,000 resamplings.

#### MFC construction and operation

A dual-chamber MFC having a total volume of 150 mL and an effective volume of 100 mL in each section (anode or cathode) was used to test the electrogenic ability of the isolated electricigen SQ-1 or *G. sulfurreducens* PCA, respectively. The anode and cathode compartments were separated by a proton exchange membrane (PEM, Nafion 117, Dupont Co., USA) with an effective area of 4π cm^2^. In the anode chamber, a carbon rod with a working area of 9.6 cm^2^ was used as an anode. The growth medium containing 20 mM sodium acetate was used as anolyte and 10% inoculum of the isolated strain or *G. sulfurreducens* PCA was added to the chamber. A small carbon brush (2.5 cm in diameter and 2.5 cm long) connected by a titanium wire served as an electrode in the cathodic chamber, which contained 50 mM potassium ferricyanide and 0.1 M PBS solution. The MFC was purged strictly with high-purity nitrogen for at least 20 min before running experiments. A 1,000 ohms resistance was connected in an external circuit except when plotting the polarization curve. Besides, the output voltages were simultaneously monitored by a Keithley instrument (Model 2400) at an interval of 24 h. The recorded voltage was converted to output power.

All the electrodes were cleaned with 5.0 M NaOH and 75% ethanol several times to remove impurities and stored in deionized water before the next use. All measurements were performed at 30 °C. The experiments were conducted in triplicate under the same operating conditions.

#### Half-cell experiments

Half-cell electrochemical experiments were performed by a 50 mL quartz bioreactor in a three-electrode system, consisting of working and counter electrodes as well as a saturated calomel reference electrode (SCE, Hg/Hg_2_Cl_2_ saturated KCl, +0.244 V vs. hydrogen standard electrode (SHE)). The polished graphite with a geometric surface area of 2.6 cm^2^ served as the working and counter electrode, respectively. The anaerobic bioreactor was sealed with a rubber plug and purged with high-purity nitrogen for 20 min before running experiments. The growth of biofilm was monitored by an 8-channel potentiostat (CHI 1040C, CH Instruments, USA) at a constant potential of 0.30 V with stirring at 30 °C. Meanwhile, the cyclic voltammogram (CV) was tested at a scan rate of 5 mV s^-1^ between the potential ranges of +0.3 V (*E*_i_) to -0.6 V (*E*_f_). All potentials in this study are versus SCE unless otherwise stated.

#### Electrochemical impedance spectroscopy

Electrochemical impedance spectroscopy (EIS) analysis was performed in a three-electrode system to evaluate the contribution of internal resistances in biofilms and this system was connected to an electrochemical station (CHI 760E, CH Instruments, USA) (Islam et al., 2017). The EIS was acquired at OCV by applying AC potential in the frequency range of 100,000-5,000,000 Hz at 10 mV amplitude to prevent biofilm detachment and maintain systematic stability. The EIS data was plotted in a Nyquist curve. The charge transfer resistance (R_ct_) and ohmic resistance (R_Ω_) were calculated by fitting the measured impedance data to an equivalent circuit (EC): R(Q[RW]) using Z view software.

#### Morphological analysis

The morphological characteristics of biofilm on carbon electrodes were analyzed with a scanning electron microscope (Hitachi S-3400N, Tokyo, Japan) at 15 kV as mentioned before (Zhou et al., 2016).

#### Analysis of ferric reduction

Ferric citrate was used as an electron acceptor to verify the capability of ferric reduction under aerobic and anaerobic conditions. The microbial growth was estimated by measuring the optical density (OD) of the cell culture at a wavelength of 600 nm (OD_600_). The *o*-phenanthroline (1,10-phenanthroline) method was used to measure the total Fe(II) concentration (Georgi et al., 2017; Islas et al., 2018). Aliquots were taken from the medium and the same volume of 0.5 M HCl was added. The mixture was vigorously shaken and incubated in an anaerobic incubator for 24 hours to extract the adsorbed Fe(II). After centrifugation at 10,000 g for 5 min, the supernatants were used for Fe(II) measurements. Experiments were performed in triplicate.

## Results and discussion

### Enrichment, isolation, and identification of the strain SQ-1

Various organic compounds were utilized in the MFC for power generation, such as glucose, sodium citrate, sodium lactate, and glutamic acid (Qiu et al., 2022; Sarmin et al., 2021; Yyya et al., 2022). The output voltage of the MFC using mixed microflora as an inoculum and sodium acetate as a carbon source was measured. It showed a maximum output voltage of 0.55 V after 35 hours of running. When there was enough sodium acetate in the culture, a voltage of 0.50 V or above was maintained for at least 50 hours in three fed-batch MFCs (Fig. 1A). Moreover, the polarization curve was plotted when the MFC stably ran in the second batch (Fig. 1B). The maximum power density was 260 mW m^-2^ when the output voltage was 0.48 V. This MFC system showed average electricity generation performance. It may be due to the antagonism between certain microbial populations or the lack of substrate metabolite-based synergy, which inhibited the biofilm enrichment on the anode or the electron transfer between microbes and electrode surface (Judith et al., 2017; Min-Chi et al., 2015).

**Figure 1.**
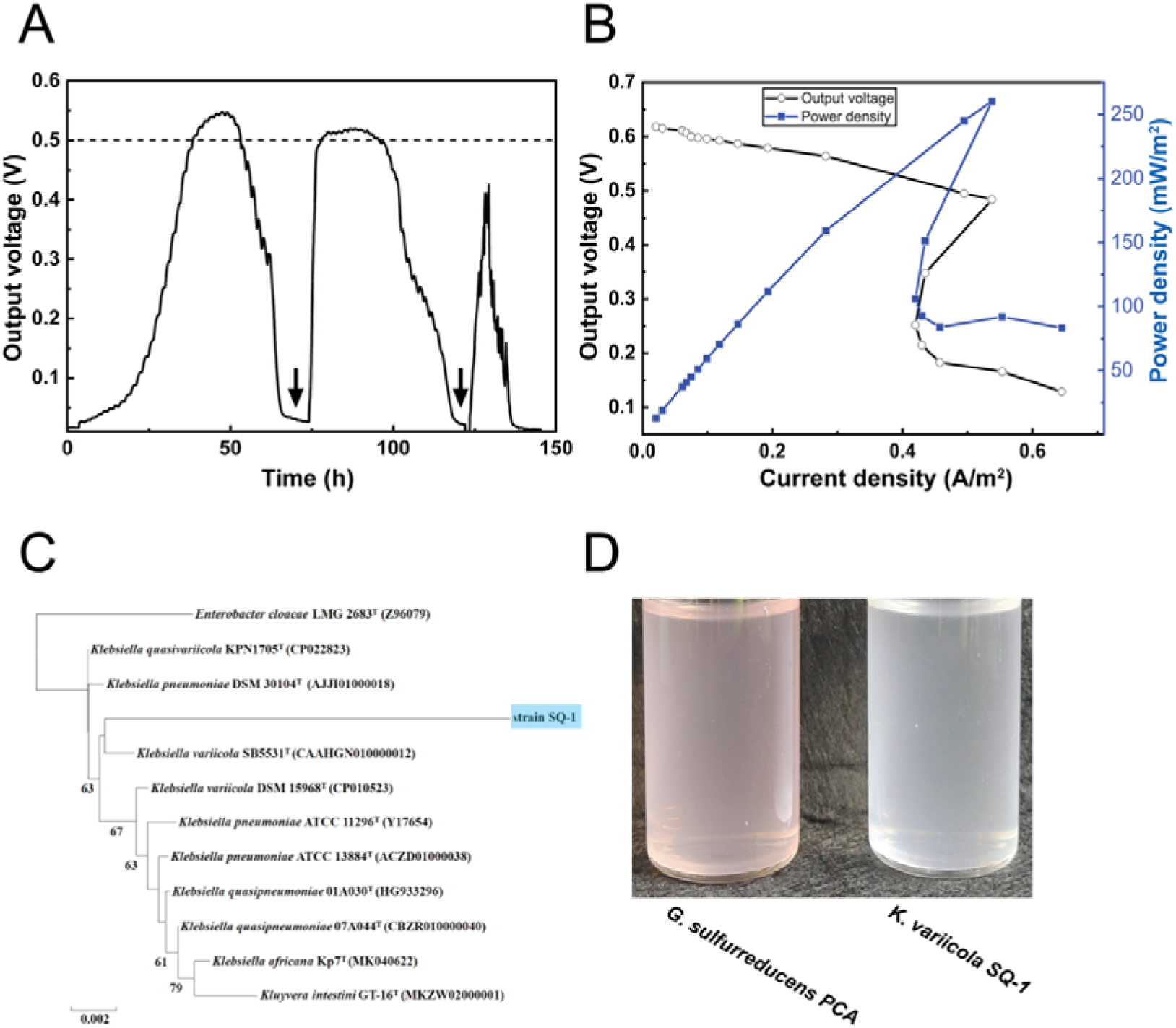
Enrichment, isolation, and identification of the strain SQ-1. (A) The output voltage performance of mixed microflora in a dual-chamber MFC using sodium acetate as a carbon source. Arrows show the supply of sodium acetate. (B) The output voltage (circle) and power density (square) curves for mixed microflora. (C) Phylogenetic analysis of the strain SQ-1 based on 16S rDNA gene sequences. (D) The appearance of the strain SQ-1 and *G. sulfurreducens* PCA in liquid growth medium after 3 days of growth.

In an MFC system catalyzed by mixed microflora, it is believed that both the biofilms enriched on the anode and free microorganisms in the solution contributed to electricity generation. Once electrochemically active biofilm was enriched from activated sludge in the MFC, the anode was used to isolate facultative anaerobic electricigens by using the spread plate technique. A single colony was picked from the agar plate and passaged five times. The isolated electricigen, named SQ-1 was identified through 16S rDNA sequencing, which was found to be 99.23% homologous to *Klebsiella variicola* SB5531^T^ (Fig. 1C). Additionally, the isolated strain SQ-1 and the strictly anaerobic electricigen *G. sulfurreducens* PCA were cultured in the same growth medium under anaerobic conditions for 3 days. Their bacterial solutions showed a different color. (Fig. 1D).

### Electricity generation performance of *Klebsiella* sp. SQ-1 in MFC

In this study, the electrochemical activity of the isolated strain SQ-1 was evaluated in MFC. The output voltage was recorded using the isolated *Klebsiella* sp. SQ-1 and *G. sulfurreducens* PCA as inoculums, respectively. The strain SQ-1 showed a rapid increase of output voltage from 110 to 180 h, which reached approximately 0.53 V (Fig. 2A). The output voltage of *G. sulfurreducens* PCA showed an identical pattern in fed-batch MFCs under similar conditions (Fig. 2B). The maximum output voltage (0.57 V) of the strain SQ-1 was higher than that of *G. sulfurreducens* PCA (0.52 V). More importantly, the strain SQ-1 was able to maintain a high output voltage for a longer time than that of *G. sulfurreducens* PCA in each fed-batch MFC, which suggested that the facultative strain SQ-1 had better electricity generation performance than either the strictly anaerobic *G. sulfurreducens* PCA or the mixed bacteria (Fig. 1A).

**Figure 2.**
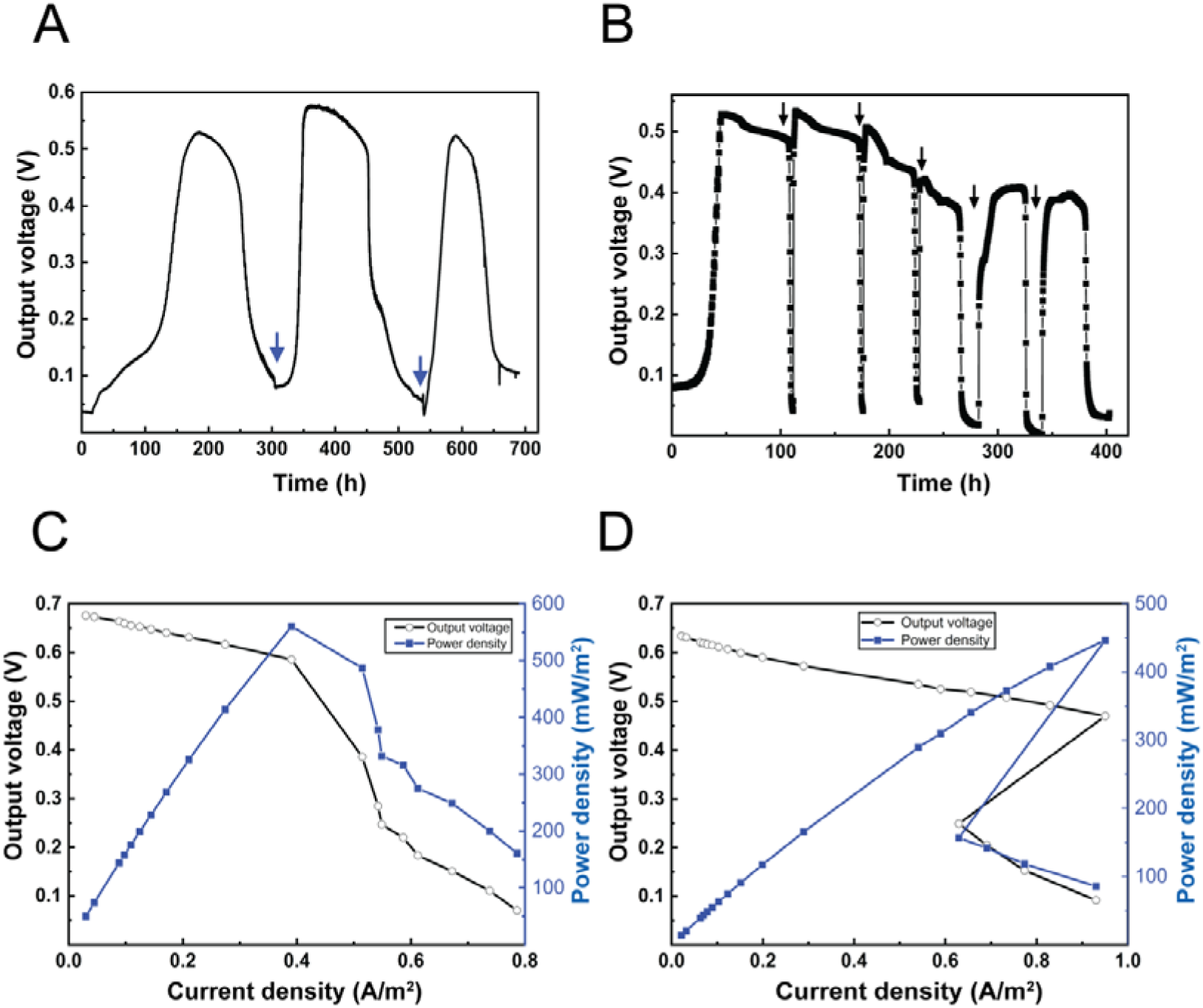
Output voltage and power density of *Klebsiella* sp. SQ-1 in MFC. (A) The output voltage performance of the strain SQ-1 in a dual-chamber MFC using sodium acetate as a carbon source. Arrows show the supply of sodium acetate. (B) The output voltage performance of a model electricigen *G. sulfurreducens* PCA in a dual-chamber MFC using sodium acetate as a carbon source. Arrows show the supply of sodium acetate. (C) The output voltage (circle) and power density (square) curve for the strain SQ-1. (D) The output voltage (circle) and power density (square) curves for *G. sulfurreducens* PCA.

The maximum power density of SQ-1 reached 560 mW m^-2^ when the corresponding output voltage was 0.59 V in MFC (Fig. 2C). The *G. sulfurreducens* PCA showed a maximum power density of 460 mW m^-2^ when the output voltage was 0.47 V (Fig. 2D). The electrical generation performance of some reported electricigens was summarized in Table 1. Facultative electricigens showed many advantages over anaerobic electricigens, for example, less stringent growth conditions and faster microbial growth. However, low power generation in a bio-electrochemical system impeded their applications in the MFC (Sunarno et al., 2019; Vikromvarasiri et al., 2016). The facultative anaerobic electricigen *Klebsiella* sp. SQ-1 with high power generation capacity might be one of the potential candidates in the engineering of biological catalysts for MFC.

**Table 1.**
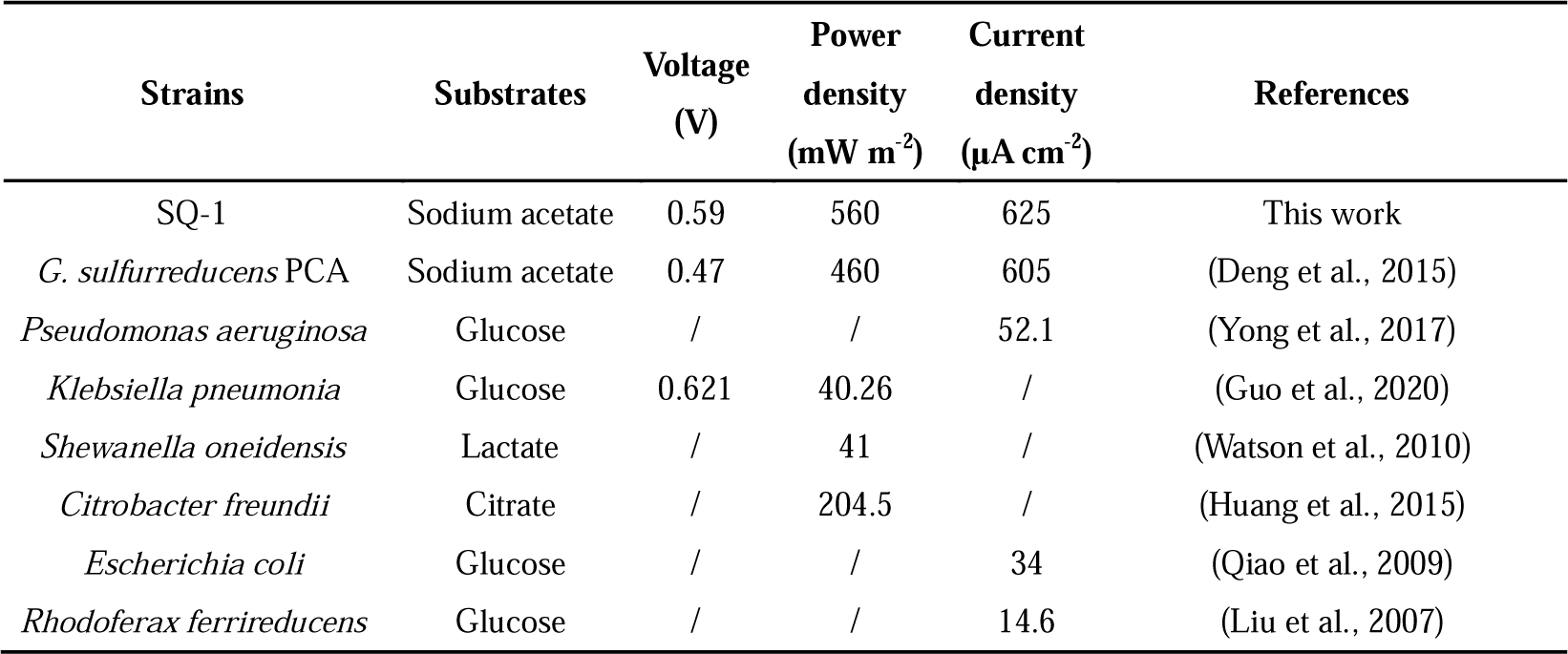
Electrical generation performance of reported 3 electricigens.

### Electrochemical performance of *Klebsiella* sp. SQ-1 in half-cell experiments

The chronoamperometric curve of SQ-1 showed that the current density increased from 7 μA cm^-2^ and drastically reached the maximum value of approximately 625 μA cm^-2^ after operating for 43 hours by utilizing sodium acetate as the sole carbon source (Fig. 3A). The power generation capacity of the strain SQ-1 was similar with that of *G. sulfurreducens* PCA as previously reported (Zhou et al., 2016).

**Figure 3.**
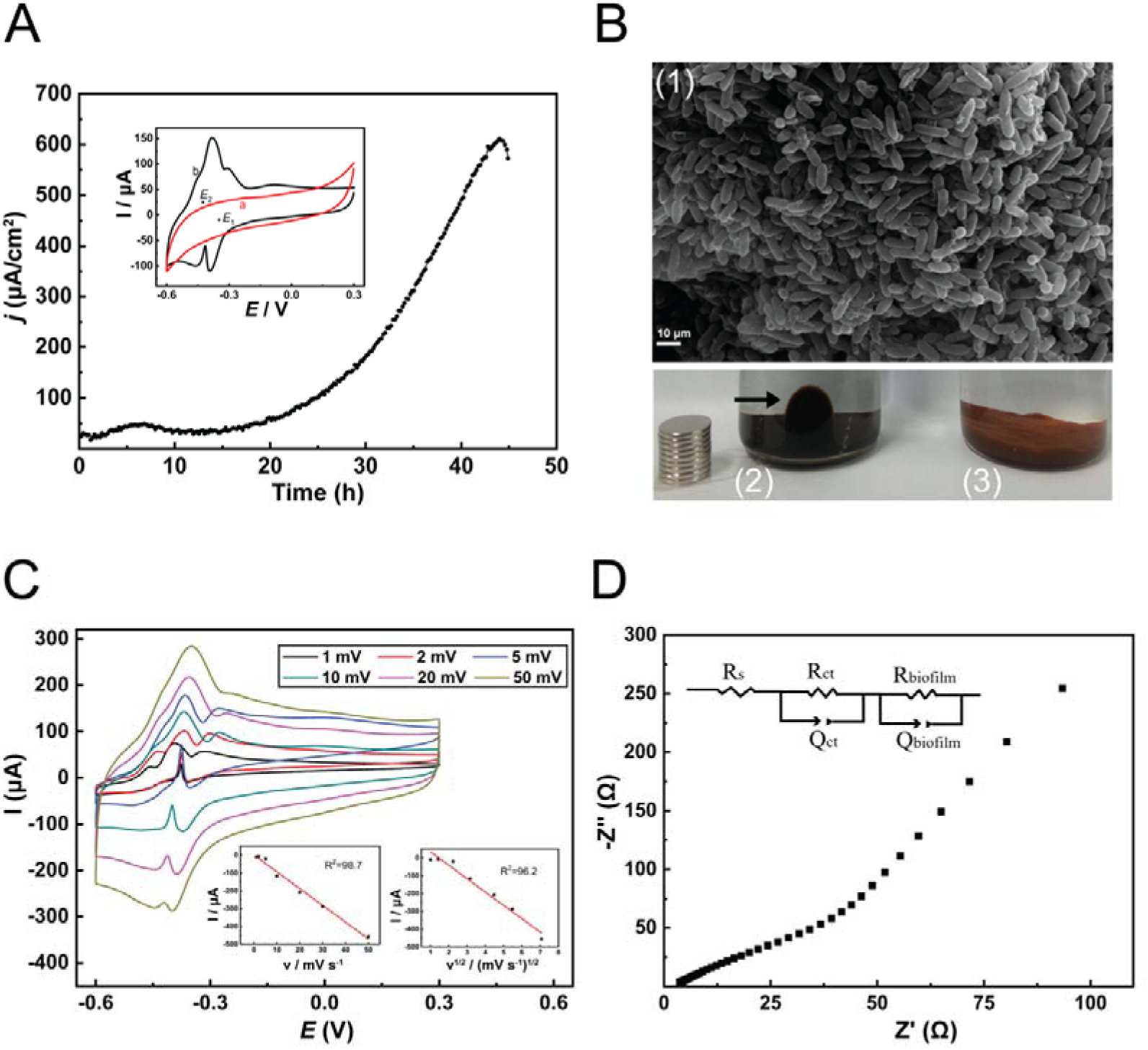
Electrochemical performance of *Klebsiella* sp. SQ-1 in a three-electrode system. (A) Chronoamperometric curves for the strain SQ-1 in a three-electrode system using sodium acetate as electron acceptors. Inset, cyclic voltammograms of the enriched strain SQ-1 biofilm on electrodes in different stages. (a) before the formation of biofilm; (b) after the formation of biofilm with a current density below 2 μA cm^-2^. (B) (B1), The morphology of the strain SQ-1 biofilms under the scanning electron microscope (5,000×). Biofilms were attached to the graphite surface. (B2) The appearance of bacterial culture after 5 days of incubation when the growth medium is supplemented with Fe(III) oxide. A magnet is placed outside the tube. The arrow shows the magnet attracts the black suspension. (B3) The initial appearance of bacterial culture. (C) Voltammograms of the strain SQ-1 biofilm. Scan rates: 1, 2, 5, 10, 20, 50 mV s^-1^. Inset, the linear dependence of the cathodic peak current density at -0.37 V versus scan rates or square roots of scan rates. (D) The Nyquist plot for biofilms attached to working electrodes.

The current density gradually decreased after it reached the plateau due to the depletion of substrate and accumulation of metabolites. The cyclic voltammogram (CV) method was performed to analyze its electrochemical characteristics, and a typical sigmoidal CV curve was obtained when the biofilm formed on a carbon sheet electrode with a maximum current density of 625 μA cm^-2^ (Table 1). The strong catalytic activity indicated the biofilms and dissociated microbes effectively transfer electrons to the electrode surface under the turnover status. Meanwhile, two pairs of redox peaks under the non-turnover status were shown (Fig. 3A inset). Their midpoint potentials were located at -0.35 V (*E_1_*) and -0.43 V (*E_2_*), respectively (Fig. 3A).

The morphological characteristics of the strain SQ-1 were studied using a scanning electron microscope (Fig. 3B(1)). Multilayered biofilms were found on the carbon electrode surface, which might show that it has enough capability to generate a highly stable current. Besides, the strain SQ-1 was inoculated into the growth medium containing Fe(III) oxide as electron acceptor and acetate sodium as electron donor, and the color of the Fe(III)oxide medium changed from brown to black after 5 days of incubation (Fig. 3B (2), (3)). The black suspension was magnetically attracted to the side of a magnet bar, indicating the strain SQ-1 has a strong ferric reduction ability.

To investigate the electron transfer kinetics of the strain SQ-1, CVs were recorded at scan rates of 1, 2, 5, 10, 20, and 50 mV s^-1^ under the non-turnover condition (Fig. 3C). A linear relationship between cathodic peak values and scan rates (ν) or the square root of scan rates (ν^1/2^) was obtained at approximately -0.37 V when the basic medium (without electron donor or acceptor) was used. Their good linear relationship (R_1_^2^=98.7 and R_2_^2^=96.3) suggested that the biofilms of SQ-1 showed surface-controlled and diffusion-controlled kinetics (Fig. 3C inset). A similar pattern of kinetics was found in *G. sulfurreducens*, *C. freundii*, or mixed culture biofilms which were formed on carbon electrodes, ITO (Indium Tin Oxide) electrodes, or gold surfaces under non-turnover conditions (Liu et al., 2015; Liu et al., 2010; Zhou et al., 2017).

Moreover, the Nyquist plot for the strain SQ-1 biofilms attached on working electrodes in a half-cell system was shown in Fig. 3D. The equivalent circuit EC: R(Q[RW]) was used to fit the impedance spectra of biofilms where Rs is the solution resistance; Rct is the charge transfer resistance at the interface of electrodes; Qct refers to the constant phase element, which represents the double layer capacitance; R_biofilm_ is the biofilm resistance; Q_biofilm_ represents the double layer capacitance caused by biofilms on the surface of electrodes. The equivalent circuit fitted well with the model and a high R_ct_ (148 ohms) was achieved when the effective electrochemical biofilms formed on the surface of electrodes.

### Ferric reduction of *Klebsiella* sp. SQ-1

The strain SQ-1 showed a typical microbial growth curve in both aerobic and anaerobic conditions when the hydrous ferric oxide (HFO) was absent (Fig. 4A). The OD_600_ value measured after 30 hours of growth in aerobic condition (OD_600_=3.46) was much higher than that in anaerobic conditions (OD_600_=0.47), which suggested that aerobic respiration plays a dominant role in the growth of strain SQ-1. To further explore the effect of HFO on the growth of *Klebsiella* sp. SQ-1, batch cultures were supplemented with different amounts of ferric citrate and incubated under aerobic or anaerobic conditions. An approximate OD_600_ value of 3.51 was measured after five days of culture in aerobic conditions without the presence of hydrous ferric oxide (Fig. 4B). A lower OD_600_ value (OD_600_=2.24) was obtained when the medium was supplemented with 5 mM HFO, indicating the growth of strain SQ-1 was affected by the extracellular HFO. However, the bacterial growth maintained at a high and similar level even if the medium was supplemented with higher concentrations of ferric citrate (10-70 mM). Therefore, it was speculated that metabolic pathways were modified in these facultative bacteria when both oxygen and HFO were present in their living environment. Aerobic respiration was the principal pathway that the strain SQ-1 utilized to grow and proliferate but a part of the energy flow was diverted to the extracellular electricigenic respiration which resulted in less biomass production. By contrast, a much lower OD_600_ value ranging from 0.1 to 0.56 was measured in anaerobic conditions (Fig. 4B) when the growth medium was supplemented with 5 mM to 70 mM ferric citrate, which suggested that the HFO served as an extracellular electron acceptor and sustained the growth of *Klebsiella* sp. SQ-1.

**Figure 4.**
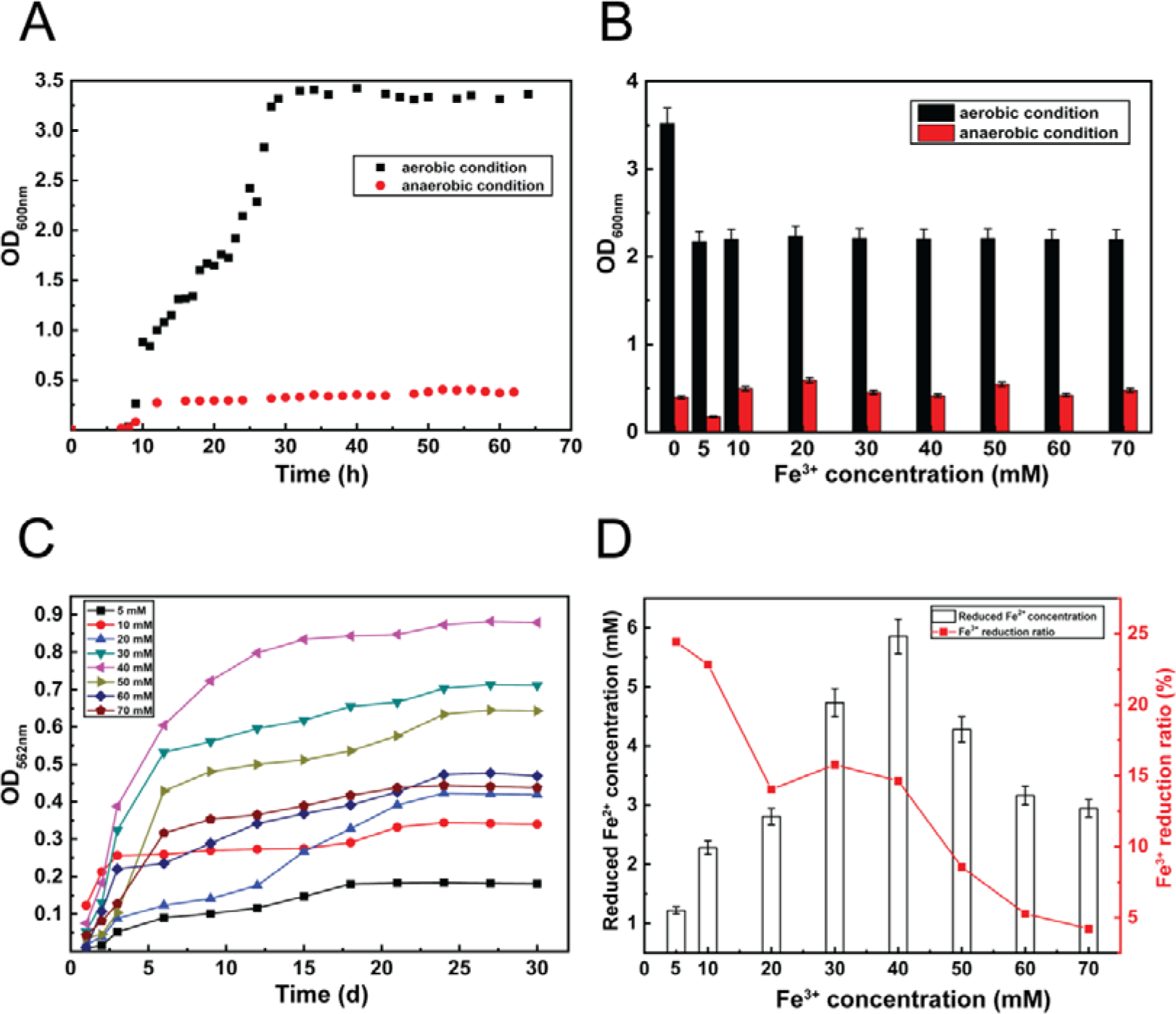
Ferric reduction of *Klebsiella* sp. SQ-1. (A) The growth curve of strain SQ-1 measured under aerobic and anaerobic conditions. (B) The growth curve of strain SQ-1 when the growth medium was supplemented with different amounts of ferric citrate. (C) Measurement of reduced Fe^2+^. (D) Analysis of Fe^3+^ reduction ratio. The concentration of Fe^2+^ was measured after 30 hours of incubation.

To estimate the Fe(III)-reduction efficiency of the strain SQ-1, the amount of reduced Fe(II) in the medium was measured by the method of *o*-phenanthroline. The amount of Fe(II) at the stationary phase gradually increased when the concentration of Fe(III) increased from 5 mM to 40mM under anaerobic conditions (Fig. 4C). The concentration of reduced Fe(II) increased from 1.25 mM to 6.00 mM (Fig. 4D). The amount of Fe(II) at the stationary phase started to fall when the concentration of Fe(III) was higher than 40 mM, which suggested that high concentration of Fe(III) might inhibit extracellular electricigenic respiration of the strain SQ-1. The highest reduction ratio of ∼25% was calculated when 5 mM Fe(III) was supplied (Fig. 4D). Notably, the bacterial biomass was at a low level (OD_600_=0.1, Fig. 4B) under this condition, which suggested that the strain SQ-1 has a strong ferric reduction ability. However, the reduction ratio decreased to approximately 15% when the concentration of Fe(III) increased to 40 mM and dropped to 4% when the concentration of Fe(III) was 70 mM (Fig. 4D).

## Conclusion

A facultative anaerobic electricigen was isolated from a wastewater tank in a biotechnology factory after a series of aerobic and anaerobic enrichment procedures. The strain SQ-1 was identified as a relative of *Klebsiella variicola* based on its 16S rDNA gene sequences. Using sodium citrate as a substrate, the maximum power density of 560 mW m^-2^ was achieved in a dual-chamber MFC, which was higher than that of *G. sulfurreducens* PCA under similar conditions. Moreover, the strain SQ-1 showed high ferric reduction activity and achieved 15% ferric reduction in the presence of 40 mM HFO under anaerobic conditions. Aerobic respiration was the optimal energy-generating way for *Klebsiella* sp. SQ-1 to grow and proliferate when cultured in aerobic conditions. When the medium was supplemented with HFO and the bacteria were cultured in anaerobic conditions, extracellular electricigenic respiration and anaerobic fermentation were employed to sustain its growth. Considering its high power generation and ferric reduction capacity, the facultative anaerobic strain *Klebsiella* sp. SQ-1 might be a promising biocatalyst in microbial electrochemistry.

## Acknowledgments

This research project was supported by grants from the Henan Province Programs for Science and Technology Development (222102110003) and the Natural Science Foundation of Henan Province (222300420259). The funding was awarded to Dongli Pei and Yayue Wang, respectively.

## Author Contributions

Lei Zhou: Data collection, Data analysis, Writing-original draft; Tuoxian Tang: Data analysis, Writing-review & editing; Dandan Deng, Writing-review; Yayue Wang, Dongli Pei, Resources, Funding acquisition, Writing-review & editing. All authors read and approved the final version of this manuscript.

